# Receptor-Based Mechanism of Cell Memory and Relative Sensing in Mammalian Signaling Networks

**DOI:** 10.1101/158774

**Authors:** Eugenia Lyashenko, Mario Niepel, Purushottam D. Dixit, Sang Kyun Lim, Peter K. Sorger, Dennis Vitkup

## Abstract

Detecting relative rather than absolute changes in external signals enables cells to make decisions in fluctuating environments and diverse biological contexts. However, how mammalian signaling networks store the memories of past stimuli and use them to compute relative signals is not well understood. Using the growth factor-activated PI3K-Akt signaling pathway, we develop computational and analytical models, and experimentally validate a novel mechanism of relative sensing in mammalian cells. This non-transcriptional mechanism relies on a new form of cellular memory, where cells effectively encode past stimulation levels in the abundance of cognate receptors on the cell surface. We show the robustness and specificity of the relative sensing for two physiologically important ligands, epidermal growth factor (EGF) and hepatocyte growth factor (HGF), and across wide ranges of background stimuli. The described memory and sensing mechanism could play a role in multiple other sensory cascades where stimulation leads to a proportional reduction in the abundance of cell surface receptors.

## Main text

Stimulation of mammalian cells with growth factors elicits a variety of context-dependent phenotypic responses, including cell migration, proliferation, and cell survival (*1*). Akt serves as a central hub in many signaling cascades activated by growth factors (*2*). Naturally, Akt phosphorylation-dependent (pAkt) pathways are implicated in multiple human diseases, such as many types of cancers (*2, 3*), diabetes (*4*) and psychiatric disorders (*5, 6*).

To understand how the immediate-early dynamics of the Akt pathway depends on the background level of growth factors, we used immunofluorescence to quantify the pAkt signal in epidermal growth factor (EGF) stimulated human non-transformed mammary epithelial MCF10A cells (SM section II). pAkt reached maximum response within minutes of continuous stimulation with EGF, and then decayed to low levels within hours (Fig. 1a). In the sensitive range of EGF concentrations, maximal pAkt response depended approximately on the logarithm of the EGF stimulus (Fig. 1b). Continuous stimulation with EGF resulted in the abundance of cell-surface EGF receptors (sEGFR) decreasing proportionally to the logarithm of the background EGF level, reaching a new steady state within hours (Fig. 1c). Prior exposure with EGF desensitized cells to subsequent EGF stimulations in a quantitative manner, i.e. maximal pAkt response to the same EGF stimulation was monotonically attenuated with increasing pre-exposure EGF levels (Fig. 1d). Thus, the pAkt response to EGF stimulation is strongly affected by background EGF levels and this effect is mediated by the removal of activated EGFRs from cell surface (*7*).

**Fig. 1:**
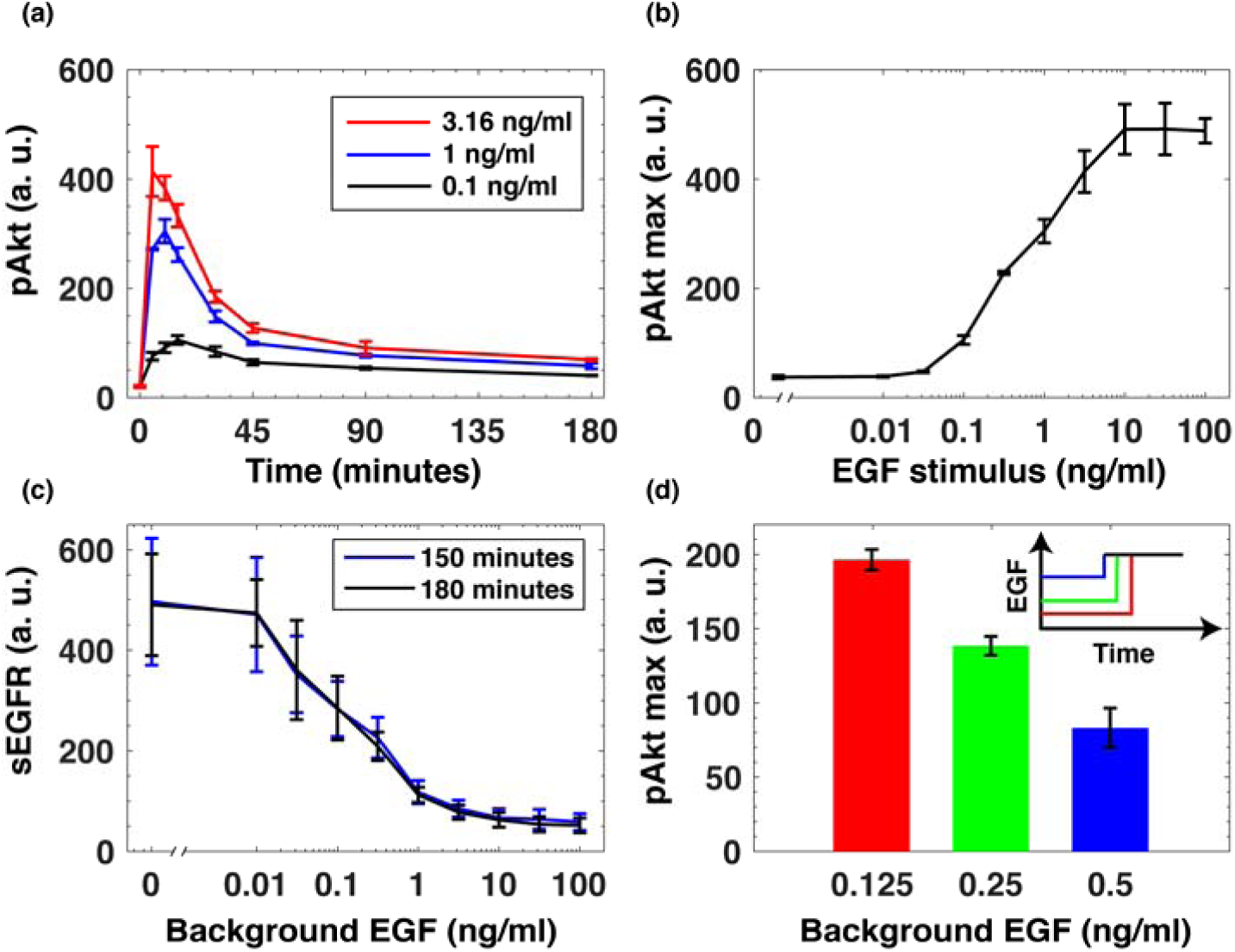
EGF-activated Akt phosphorylation and desensitization. (a) Time-dependent levels of phosphorylated Akt (pAkt) in MCF10A cells exposed to increasing stimulation with extracellular EGF. (b) Maximal pAkt response as a function of EGF stimulus. (c) Steady state levels of surface EGFR (sEGFR) after 150 and 180 minutes of stimulation with a constant dose of EGF. (d) Desensitization of the maximal pAkt response. In three different experiments (represented by bars and inset curves with different colors) MCF10A cells were pre-treated with increasing background doses of EGF for three hours, followed by a second stimulation with 2 ng/ml of EGF. Error bars represent the standard deviation of technical replicates.

To understand how background EGF levels affect the pAkt response to subsequent EGF stimulation we constructed an ordinary differential equation (ODE) model of EGF-dependent Akt phosphorylation. The model included several well-established features of the EGFR signaling cascade (ref. (*8*), SM section III) (Fig. 2a). We constrained the ranges of model parameters based on literature-derived estimates (SM Table 1), and fitted the model using pAkt time courses and steady state sEGFR levels at different doses of EGF stimulations. We used simulated annealing to optimize model parameters (SM section III), and considered multiple distinct parameter sets from the optimization runs for further computational analysis (SM Fig. 9a, b).

**Fig. 2:**
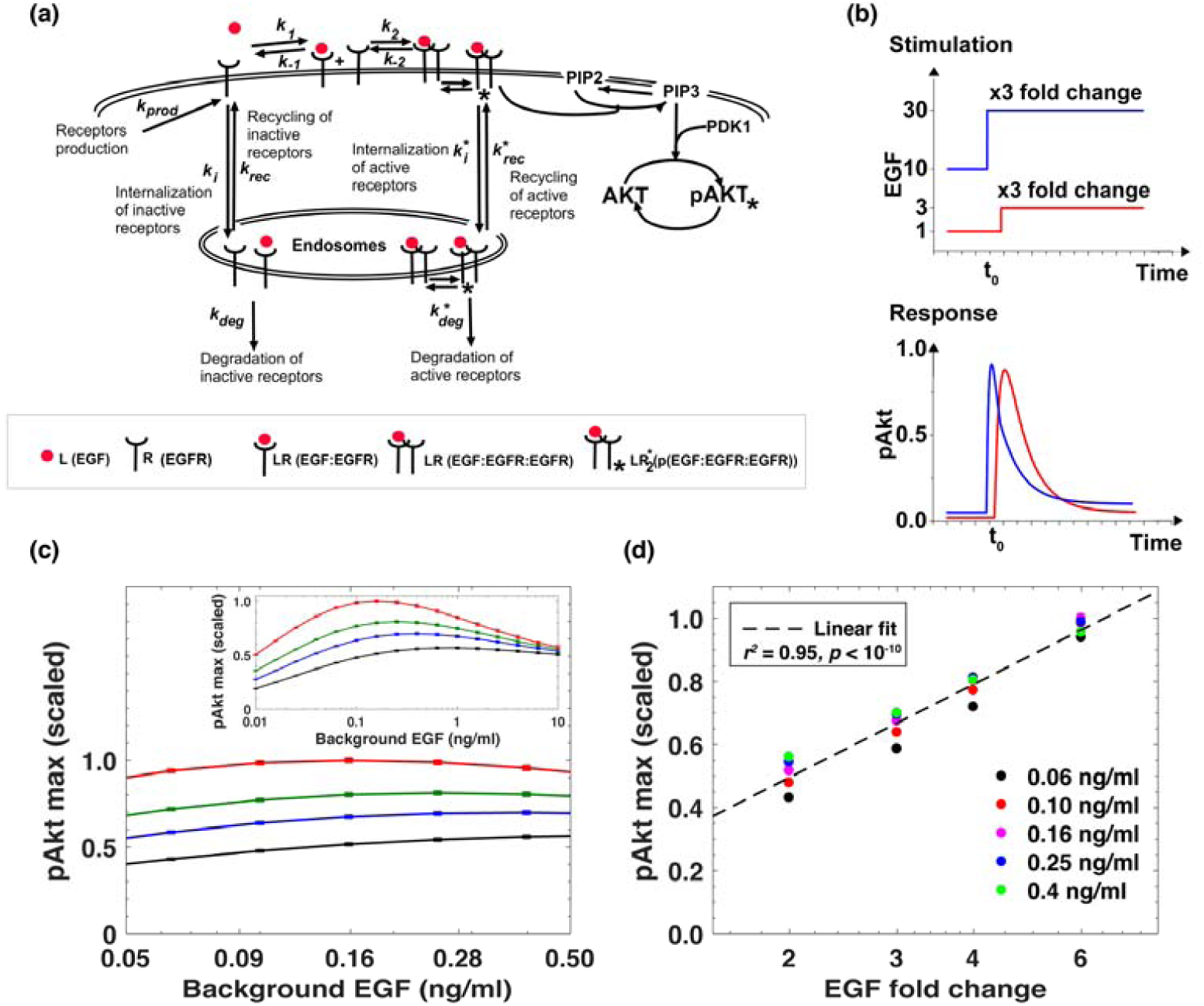
Computational model predicts relative sensing of EGF levels. (a) Computational model of EGFR signaling leading to phosphorylation of Akt. Rate constants with asterisks refer to reactions associated with activated (phosphorylated) receptors. Only a subset of reactions in the network are shown for brevity. (b) Protocol used to explore relative sensing, showing EGF stimulation (top) and the corresponding pAkt response (bottom). Cells were first exposed to different background EGF stimulations (blue and red), and were next subjected to the same fold change in EGF at time *t*_*0*_. The resulting maximal pAkt responses were similar, indicating relative sensing. (c) The maximal pAkt response observed after exposing the ODE model *in silico* to different background EGF levels (x axis), followed by a 2-, 3-, 4-, or 6-fold increase (different colors) of EGF; inset shows pAkt response over a wider range of background EGF levels. (d) Maximum pAkt responses from stimulations with various EGF background levels (indicated by data points with the same shape and color) were combined and plotted as a function of the fold change in EGF dose (x axis). Dashed line represents log-linear fit to data (Pearson’s *r*^*2*^ = 0.96, *p* < 10^−15^). Error bars represent the standard deviation of top 10 model fits.

Using the fitted dynamical model, we explored the ability of the Akt pathway to respond to relative, rather than absolute, changes in EGF levels. We simulated the pAkt response by exposing the model *in silico* to a range of background EGF levels. We then simulated different fold change increases in EGF concentration (Fig. 2b). The model predicted that the maximal pAkt response indeed depends primarily on the EGF fold change relative to the background stimulation (Fig. 2c). Relative sensing occurred over an order of magnitude of background EGF concentrations and the resulting pAkt response was approximately proportional to the logarithm of the EGF fold change (Fig. 2d). Notably, the model predicted relative sensing in the range of EGF background concentrations where endocytosis was sensitive to background ligand stimulation. At low EGF background concentrations (< 0.01 ng/ml), no substantial sEGFR removal was predicted at steady state (SM Fig. 9b), and consequently there was no significant desensitization of the pAkt response. In that regime the pAkt response after a step change depended primarily on the absolute EGF level. On the other hand, at high background EGF concentrations (> 1 ng/ml), a large fraction of sEGFR was removed from cell surface and consequently the network responded only weakly to further EGF stimulation.

We next experimentally tested the model-predicted relative sensing behavior in MCF10A cells. Cells were first treated with various background EGF concentrations for three hours to ensure steady state sEGFR levels (Fig. 1d), and that pAkt had decayed after a transient increase (Fig. 1c). As in the computational analysis (Fig. 2b), cells were then exposed to different fold changes in EGF levels. pAkt levels were measured at 2.5, 5, 10, 15, 30 and 45 minutes after the step increase in EGF stimulation (SM Fig. 1a, b); similar results were obtained in two independent biological replicates (SM Figures 3 and 4a, b). The experiments confirmed the predictions of the computational model that maximum pAkt response depends primarily on the fold change in EGF levels and not its absolute concentration (Fig. 3a, SM Fig. 2). Specifically, across more than an order of magnitude of EGF background concentrations (0.03 - 0.5 ng/ml) the same EGF fold change (lines of the same colors in Figure 3a) elicited similar pAkt responses. The concentration range in which relative sensing was observed is consistent with recent estimations of *in vivo* EGF levels (*9*). Finally, in good agreement with model predictions, the maximum pAkt response was approximately proportional to the logarithm of EGF fold change (Fig. 3b).

**Fig. 3:**
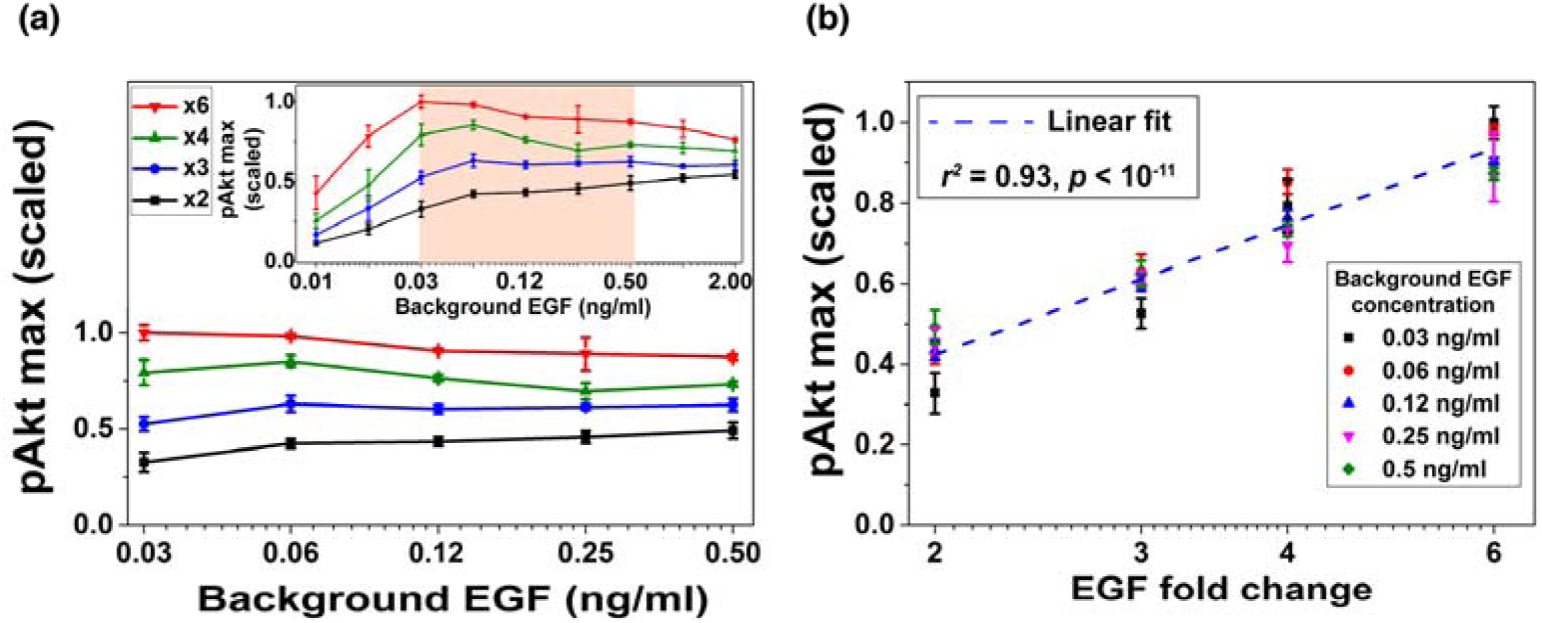
Experimental confirmation of relative sensing of extracellular EGF levels by pAkt. (a) The maximum pAkt responses after exposing MCF10A cells to different background EGF levels (x axis) for 3 hours followed by 2-, 3-, 4-, and 6-fold increases (different colors) in EGF. Inset shows experimental pAkt response over a wider range of background EGF levels. (b) Maximum pAkt responses to fold changes in EGF depended approximately logarithmically on the fold change. Maximum pAkt responses from experiments with various EGF background levels (indicated by data points with the same shape and color) were combined and plotted as a function of the fold change in EGF dose (x axis). Dashed line represents log-linear fit to the data (Pearson’s *r*^*2*^ = 0.93, *p* < 10^−11^). Error bars represent the standard deviation of technical replicates.

**Fig. 4:**
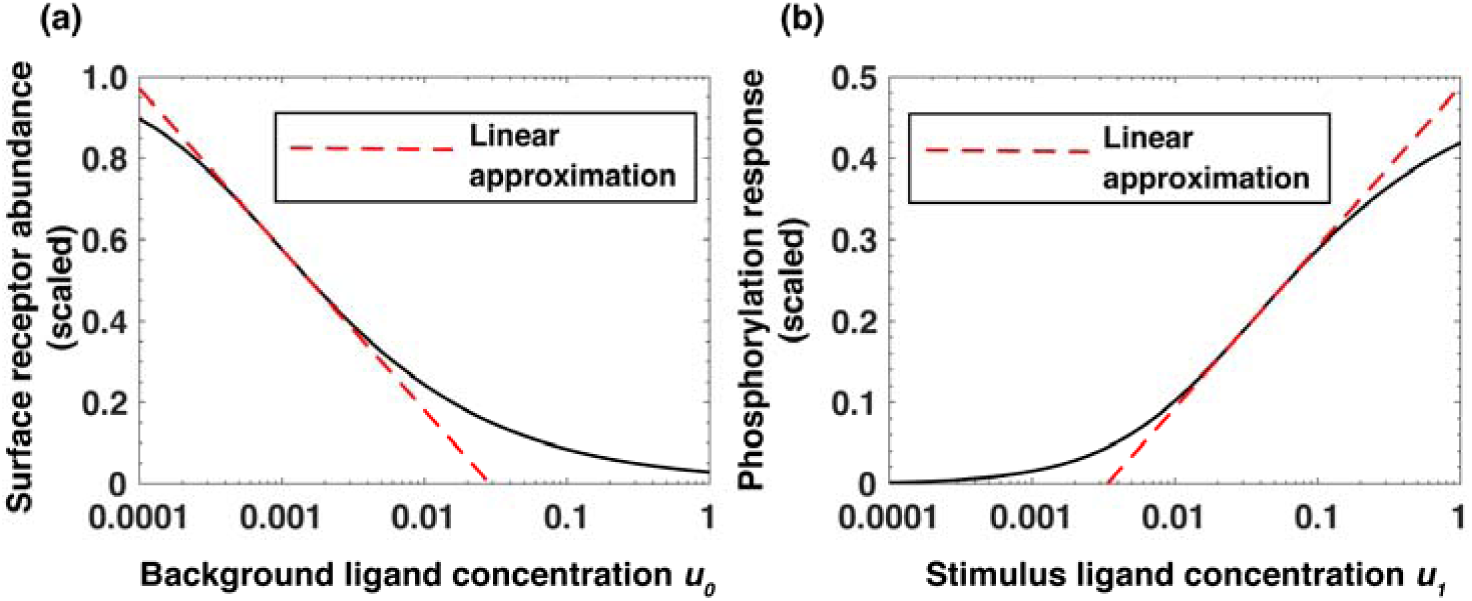
An analytical model of receptor-based memory and relative sensing. (a) Approximate log-linear dependence of the scaled steady-state receptor abundance [*R*]_*T*_ on the normalized background ligand concentration *u*_*0*_ = [*L*]_0_/*K*_*d1*_, where *K*_*d1*_ is the equilibrium dissociation constant of EGF binding to EGFR. (b) Approximate log-linear dependence of the maximum phosphorylation response on the normalized ligand stimulus *u*_*1*_ *=*[*L*]_1_/*K*_*d1*_. Dashed red lines represent the log-linear approximation.

To better understand the mechanism responsible for the observed relative sensing of extracellular EGF concentration, we constructed a simplified analytical model of the signaling network (SM section IV). This model revealed that, across a broad range of background concentrations, the steady-state abundance of cell surface receptors [*R*]_*T*_ decreases approximately log-linearly as a function of the background ligand (EGF) concentration [*L*]_0_ (Equation 1 and Fig. 4a):

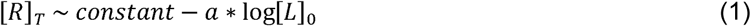

and that the maximal receptor phosphorylation response [*LR*^*\**^_2_] depends approximately log-linearly on the level of the subsequent stimulation [*L*]_1_ and linearly on the steady-state receptor abundance [*R*]_*T*_ (Equation 2, Fig. 4b):

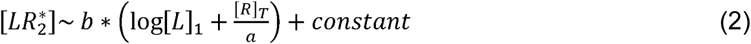

where *a* and *b* are numerical constants (SM section IV). As a result of these relationships, the phosphorylation response [*LR*^*\**^_2_] after an increase in ligand concentration from [*L*]_0_ to [*L*]_1_ depends, in agreement with computational and experimental analyses, approximately on the logarithm of the stimulation fold change [*L*]_1_/[*L*]_0_ (SM section IV):

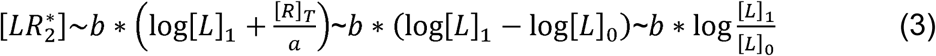

The analytical model revealed that the range of the background ligand concentrations where the relative sensing is observed is primarily determined by two aggregate parameters. One quantifying the ability of the circuit to capture the EGF signal and elicit a phosphorylation response (Equation 24a, section IV of SM). The other quantifying the ability of the circuit to preferentially internalize and degrade active (phosphorylated) receptors compared to inactive receptors (Equation 24b, section IV of SM). The analysis of the model also demonstrated that the relative sensing and receptor-based memory is robust to perturbations in network parameters (SM section IV).

In addition to EGF, Akt phosphorylation can be induced by multiple other ligands, including hepatocyte growth factor (HGF) (*10*) which binds to its cognate receptor cMet (*11*). To investigate the specificity of the receptor-based cell memory to past ligand exposures, we used the two ligands, EGF and HGF, which share most of their signaling components downstream of their cognate receptors (*12*). We stimulated cells with background doses of either HGF or EGF for three hours, and then treated cells using either the same or the other growth factor to elicit pAkt response (Fig. 5a, b). Pre-exposure with HGF did not substantially downregulate EGF-induced pAkt responses, but substantially decreased HGF-induced responses (Fig. 5a). Similarly, there was a relatively small desensitization of HGF-induced responses due to pre-exposure with EGF, while a significant desensitization of EGF-induced pAkt responses was observed (Fig. 5b). We further confirmed that exposure of MCF10A cells to various concentrations of HGF leads to pronounced HGF-dependent removal of cMet from the cell surface, without significant removal of sEGFR (SM Fig. 5a). Similarly, the pre-exposure of cells to EGF leads to EGF-dependent removal of sEGFR without a significant change in surface cMet abundance (SM Fig. 5b). These observations support the mechanism in which the relative sensing of extracellular ligands relies on the memory of their past exposures encoded primarily in the abundances of their cognate cell-surface receptors.

**Fig. 5:**
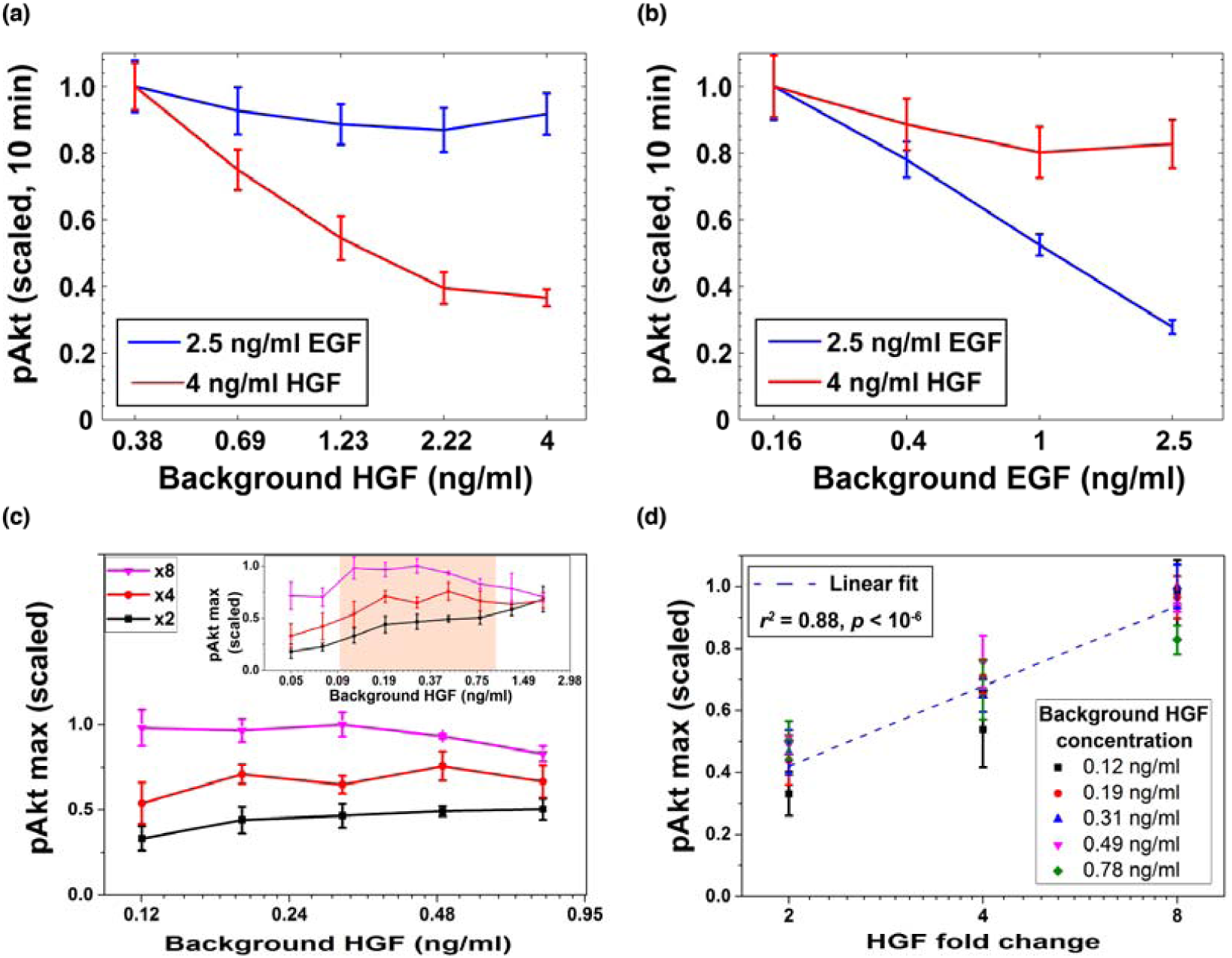
Desensitization and cell memory for EGF- and HGF-induced pAkt response. MCF10A cells were exposed to various background doses of either HGF or EGF for three hours, and then stimulated using either the same or the other growth factor. pAkt levels were then measured 10 minutes after the addition of the second stimulus. (a) EGF (blue, 2.5 ng/ml) or HGF (red, 4 ng/ml) induced pAkt response in cells pre-exposed with various doses of HGF (shown on the x axis) for three hours. (b) EGF (blue, 2.5 ng/ml) or HGF (red, 4 ng/ml) induced pAkt response in cells pre-exposed with various doses of EGF for three hours. Error bars represent the standard deviation of technical replicates. (c) The maximum pAkt response in MCF10A cells exposed to different background doses of HGF (x axis) for 3 hours followed by 2-, 4-, and 8-fold increase (different colors) of HGF. Inset shows experimental pAkt response over a wider range of background HGF levels. (d) The maximum pAkt responses to HGF fold changes depended approximately logarithmically on the fold change. Maximum pAkt responses from experiments with various HGF background levels (indicated by data points with the same shape and color) were combined and plotted as a function of the fold change in HGF dose (x axis). Dashed line represents log-linear fit to data (Pearson’s *r*^*2*^ = 0.88, *p* < 10^−6^). Error bars represent the standard deviation of technical replicates.

Given the observed HGF-dependent removal of cell surface cMet receptors and resulting pAkt desensitization, we investigated whether the maximum pAkt response depends, similarly to EGF, on the relative fold changes in the level of extracellular HGF. To that end, we exposed cells to a range of different background levels of HGF, and then stimulated cells with different fold changes in HGF concentrations (Fig. 5c,d and SM Fig. 6). These experiments demonstrated that HGF-induced phosphorylation of Akt also depends primarily on the fold change in extracellular HGF concentration across almost an order of magnitude of background HGF exposures (between 0.1 and 1 ng/ml HGF) and can distinguish up to 8 fold changes in HGF concentration (Fig. 5c). Moreover, similar to EGF, the maximum pAkt levels depends approximately log-linearly on the fold change in HGF (Fig. 5d).

**Fig. 6:**
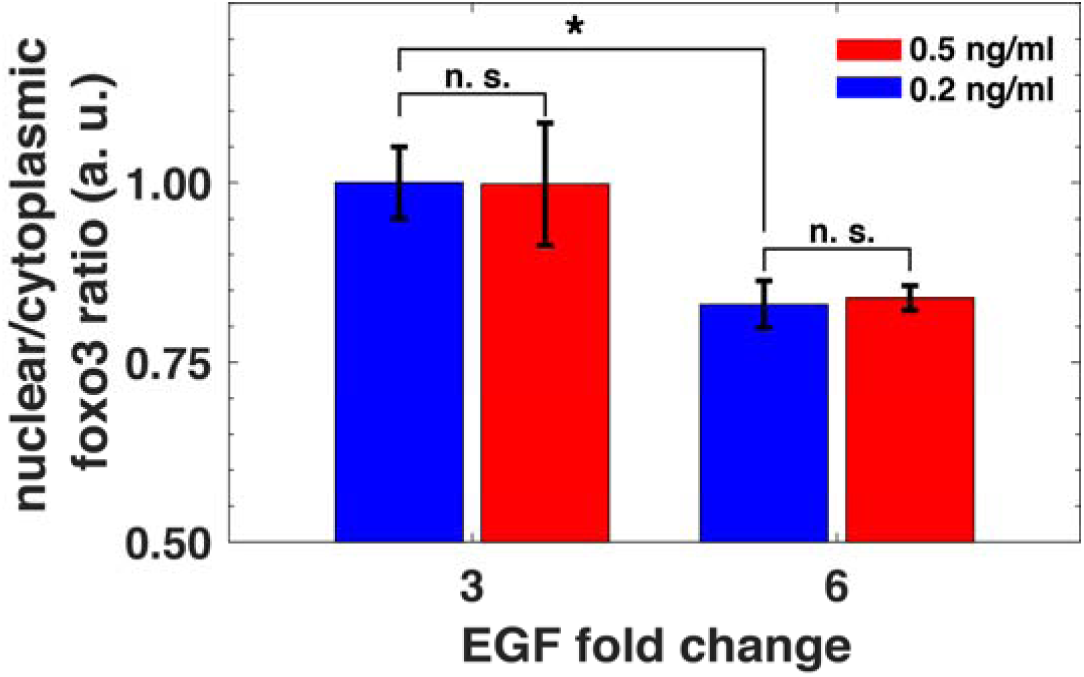
Relative sensing of EGF concentrations by FoxO3. MCF10A cells were exposed to two background doses of EGF for three hours, and then treated with 3- and 6-fold increase in EGF concentrations. The ratio of nuclear-to-cytoplasmic FoxO3 levels (y-axis) was measured using quantitative immunofluorescence (see SM section II) after 15 minutes of the EGF fold changes. Following the patterns observed for pAkt activation, the ratio of nuclear-to-cytoplasmic FoxO3 level depended on the relative but not absolute change in EGF concentration. Statistical significance was calculated using the Wilcoxon rank sum test (n = 5); * corresponds to p < 0.01, and n. s. corresponds to p > 0.1. Error bars represent the standard deviation of technical replicates.

Relative sensing of extracellular ligands should affect important biological targets of the PI3K-Akt pathway. The FoxO3 transcription factor is a key effector of the pathway, and is involved in multiple cellular processes including apoptosis, proliferation, and metabolism (*13*). Akt phosphorylation of FoxO3 leads to its translocation from the nucleus to cytoplasm and subsequent transcriptional deactivation (*13*). To investigate FoxO3 activity following EGF stimulation, we used quantitative immunofluorescence to measure its nuclear-to-cytoplasm ratio (*14*). We exposed cells to two different background EGF levels for three hours, and then treated them with two different fold changes in EGF concentrations. Consistent with relative sensing by pAkt, the nuclear-to-cytoplasmic ratio of FoxO3 reflected the relative, but not the absolute changes in EGF stimulation (Fig. 6 and SM Fig. 8). Thus, relative sensing of the signal is faithfully transmitted to at least some of the physiologically important effectors of the PI3K-Akt pathway.

The non-transcriptional receptor-based mechanism of cell memory and relative sensing described in our paper operates on the time scales of several minutes to hours. We primarily used the maximal level of pAkt to demonstrate the relative sensing capabilities of the network. However, relative sensing is also observed for the time integral of pAkt levels (SM Fig. 7). The developed analytical model further reveals that relative sensing does not depend on receptor dimerization, and that a similar mechanism can function in a pathway where a signal is initiated by monomeric receptors (SM section IV).

Receptor endocytosis and down-regulation following ligand stimulation has been canonically associated with signal desensitization (*15-17*). Our analysis suggests a specific quantitative role for receptors downregulation: it may allow cells to continuously monitor signals in their environment (*18-20*) and respond to relative changes in environmental stimuli. Recent elegant studies (*21-23*) have demonstrated that transcriptional motifs may efficiently buffer cell-to-cell variability in signaling components when responding to a constant stimulation. In contrast, our study describes a non-transcriptional mechanism of sensing changes relative to past extracellular stimulation. These two processes are likely to be complementary, thus allowing cells to sense, across different timescales, relative changes in environmental signals while also buffering cellular variability (*24, 25*).

Although there are usually ∼10^5^-10^6^ EGFR receptors on mammalian cell surface (*26*), the downstream network response, for example Akt phosphorylation, often saturates when only a relatively small fraction (5-10%) of the receptors are bound to their cognate ligands (*8, 26*). Our study suggests that one potential advantage of such a system architecture is that, beyond simple signal activation, it may endow cells with a large dynamic range of receptor abundances to memorize stimulation levels of multiple extracellular ligands (*27, 28*). Signal-mediated removal has been reported for many other receptors and membrane proteins, such as the G protein coupled receptors (GPCRs) (*29*) involved in multiple sensory systems and AMPA-type glutamate receptors (*30*) implicated in synaptic plasticity. Therefore, similar relative sensing mechanisms may be important in multiple other receptor-based signaling cascades and different biological contexts.

